# Signatures of neutral evolution in exponentially growing tumors: a theoretical perspective

**DOI:** 10.1101/2020.07.29.227454

**Authors:** Hwai-Ray Tung, Rick Durrett

**Affiliations:** Dept. of Math, Duke University

## Abstract

We investigate the site frequency spectrum in the two-type model of clonal evolution. If the fitnesses of the two types are λ_0_ < λ_1_ then the site frequency spectrum is *c*/*f^α^* where *α* = λ_0_/λ_1_. This is due to the advantageous mutations that produce the founders of the type 1 population. Mutations within the growing type 0 and type 1 populations follow the 1/*f* law. Our results show that neutral evolution can be distinguished from the two-type model using the site frequency spectrum.

## 1 Introduction

Several years ago, Sottoriva and Graham [11] made the controversial claim that the 1/*f* site frequency spectrum that occurs in neutrally evolving homogeneously mixing populations of constant size, fits mutant allele frequencies in next generation sequencing of tumor bulk samples. In work published in Nature Genetics [14], they show that 323 of 904 samples from 14 cancer types showed excellent straight line fits when the cumulative number of mutations of frequency ≥ *f* is plotted versus 1/*f*, see Figure 2b in [14].

This paper has been cited 200 times, but among these works there are a number of papers criticizing the result. See [10, 13, 3]. The December 2018 issue of Nature Genetics contains three letters raising objections to the conclusion [1, 12, 8]. Four common criticisms are

i. Inferring the allele frequency *f* requires accurate estimates of local copy number and ploidy. In addition, Wu et al [13] point out that local samples may not be indicative of overall frequencies.
ii. Failure to reject the null model is not the same as proving it is true. To quote MacDonald, Chakrabarti, and Michor [8] “The fact that a model of neutral evolution leads to a linear relationship between *M*(*f*) (the number of mutations with frequency ≥ *f*) and 1/*f* does not imply … the presence of neutral evolution.”
iii. Tarabachi et al [12] applied methods that look at the *dN/dS* ratio, which compares the number of nonsynonymous and synonymous mutations, to look for signs of selection. They claim to have found significant signs of selection in tumors that were classified as neutral. However when the analysis was repeated on publicly available pancreatic cancer data, Graham, Sottoriva et al found no values significantly different from 1.
iv. Tarabachi et al [12] say “the deterministic models of tumor growth described by Williams et al [14] rely on strong biological assumptions. Using simple branching process to simulate neutral and nonneutral growth, they show that *R*^2^ > 0.98 is neither necessary nor sufficient for neutral evolution.”

To try to shed some light on the controversy, we will compute the site frequency spectrum produced by the two-type model of clonal evolution. We will describe the model in Section 3. The two-type model and its m-type generalization have been extensively studied. See [5] for results and references. This model is relevant to the discussion of [14] because it appears in the criticisms of MacDonald, Chakrabarti, and Michor [8] and Bozic, Patterson, and Waclaw [3].

## 2 Simple derivations of the 1/*f* spectrum

Sottoriva and Graham say in their original paper [11] that “the power law signature is common to multiple tumor types and is a consequence of the effectively-neutral evolutionary dynamics that underpin the evolution of a large proportion of cancers.” To explain the source of the 1/*f* curve in an exponentially growing tumor, we give the derivation of the 1/*f* frequency distribution from [14]. They assumed that cells divide at rate λ and use *N*(*t*) to be the number of cells at time t. If we assume that the mutation rate is μ (which we assume takes into account their ploidy parameter *π*), then the expected number of new mutations before time *t*, *M*(*t*), satisfies

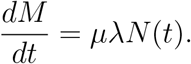

Solving gives

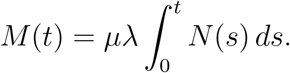

Since *N*(*s*) = *e*^λ*s*^ (we have absorbed their parameter *β* into λ), we observe that a mutation that occurs at time *s* will have frequency *e*^−λ*s*^ in the population and that the prior expression for *M*(*t*) can be simplified to

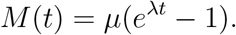

Ignoring the −1, if we set *t* = −(1/λ) log *f* to make the frequency *f*, then

#### Theorem 1.

*The number of mutations with frequency* ≥ *f is*

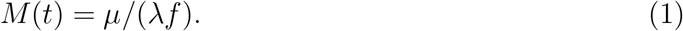

From the derivation given above, we see that the 1/*f* site frequency spectrum comes from the fact that mutations occur at a rate proportional to the size of the population and the fact that the population is growing exponentially fast.

As the authors acknowledge in [14], the site frequency spectrum in an exponentially growing branching process was derived earlier in [4]. To formulate their result, consider a continuous time branching process in which individuals give birth at rate *a*_0_ and die at rate *b*_0_ < *a*_0_. Let λ_0_ = *a*_0_ − *b*_0_ be the exponential growth rate and *γ* = λ_0_/*a*_0_ be the probability that the branching process starting from a single individual does not die out. For simplicity it is assumed that mutations occur continuously during the lifetime of the cell, rather than only at birth. This has very little effect on the behavior of the model, but it does change the constant.

Let *F*_*t*_(*x*) be the expected number of neutral mutations that are present in a fraction ≥ *x* of the individuals at time *t*.

### Theorem 2.

*If neutral mutations occur at rate v, then when the branching process does not die out*

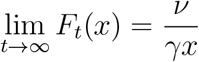

The 1/λ disappears from (1) since births occur throughout the life of a cell rather than only at birth. The 1/γ comes from the fact that if *Y*_0_(*t*) is the number of individuals at time t that have an infinite line of descent, then *Y*_0_(*t*)/*Z*_0_(*t*) → γ. In the set-up of Theorem 1, no cells die so γ = 1. The mutations occurring to the *Z*_0_(*s*) − *Y*_0_(*s*) individuals whose family line dies out will be lost from the population unless *s* is close to t.

## 3 A two-type model

McDonald, Chakrabarti, and Michor [8] consider two alternative evolutionary models in order to argue that other underlying models can produce a linear relationship between 1*/f* and the cumulative number of mutations with frequency ≥ *f*. Their second model is an infinite alleles branching process model previously studied by McDonald and Kimmel [9]. We will ignore this model, since in studying DNA sequence data the appropriate mutation scheme is the infinite sites model.

In their first model, clonal expansion begins with a single cell of the original tumor-initiating type (type 0). To keep the notation consistent with [4] and [5], we will suppose that type 0 individuals give birth at rate *a*_0_ and die at rate *b*_0_, so the exponential growth rate is λ_0_ = *a*_0_ − *b*_0_. For simplicity, we will suppose that neutral mutations accumulate during the individual’s life time at rate v, instead of only at birth.

Type 0 individuals mutate to type 1 at rate *u*_1_. Type 1 individuals give birth at rate *a*_1_ and die at rate *b*_1_. Their exponential growth rate is λ_1_ = *a*_1_ − *b*_1_ where λ_1_ > λ_0_. In [8], different type 1 families have different increases in their growth rates that follow a normal distribution. In this section, we will assume all type 1 mutations have the same growth rate. In Section 4, we will consider the implications of random fitness changes for the behavior of the model.

As in [8], we will, for simplicity, restrict our attention to two types. The type 0’s are a simple branching process, so well-known results show that

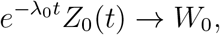

where *W*_0_ = 0 with probability *b*_0_/*a*_0_ and has a rate λ_0_/*a*_0_ exponential distribution with probability λ_0_/*a*_0_.

The reader will see many complicated formulas in this paper, so it will be useful to have a concrete set of parameters to plug into these formulas. Borrowing an example from [5], we will set

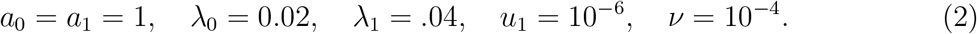

We do not pretend that these parameters apply to any specific cancer, but for a mental picture, you can imagine that type 0s are colon cancer cells in which both copies of APC have been knocked out, while type 1 cells in addition have a KRAS mutation.

At time t = 1000, *e*^λ_0_*t*^ = *e*^20^ = 4.852 × 10^8^. To account for *W*_0_ we note that the mean of (*W*_0_|*W*_0_ > 0) is *a*_0_/λ_0_ = 50, so the population size at time 1000 is 2.43 × 10^10^ times an exponential(1) random variable. A 1 cm^3^ tumor has about 10^9^ cells. (24.2)^1/3^ = 2.90, so the tumor is roughly a cube 3 cm on a side.

The study of the second wave is simpler if we suppose that 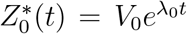 for all *t* ∈ (−∞, ∞). Mutations from type 0 to 1 occur at rate *u*_1_. Let *σ*_1_ be the time of the first successful type 1 mutation, i.e., one whose branching process does not die out. Durrett and Moseley [7] showed, see (29) in [5], that *σ*_1_ has median

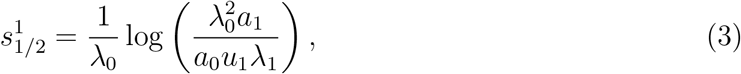

and as *u*_1_ → 0

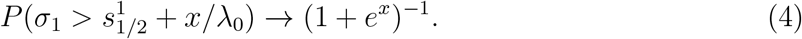

In the concrete example

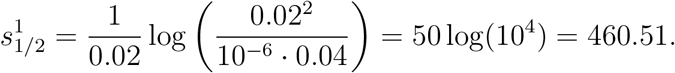

In colon cancer where cells divide every four days, 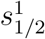 is 1842 days or a little more than 5 years.

Durrett and Moseley were the first to rigorously prove results about the asymptotic behavior of the size of the type 1 population 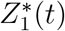, see Section 9 of [5]. Durrett [4] noticed that the constants are simpler if we use a different normalization.

### Theorem 3.

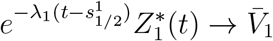. *The Laplace transform of* 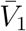 *is*

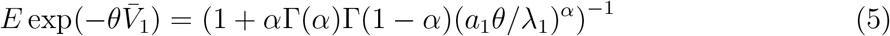

*where* 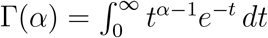.

Replacing *θ* by λ_1_*θ*/*a*_1_ in (5) we see that the distribution of 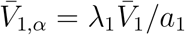 depends only on α. In our concrete example *α* = 1/2.

To prove results about the site frequency spectrum, we have to delve into the details of the proof. A mutation that occurs at time *s_i_* makes a contribution of 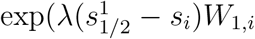 to the limit in Theorem 3 where

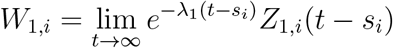

and *Z*_1.*i*_ is the type 1 branching process initiated by the mutation at time *s_i_*. *W*_1,*i*_ = 0 with probability *b*_1_/*a*_1_ and has an exponential(λ_1_/*a*_1_) distribution with probability λ_1_/*a*_1_. The points (*s_i_*, *W*_1,*i*_) with *W*_1,*i*_ > 0 are a Poisson process on (−∞, ∞) × (0, ∞) with mean measure

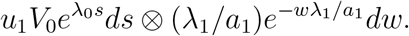

If we map these points

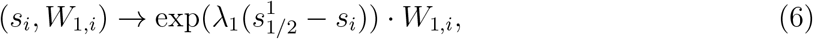

then we get a Poisson process on (0, ∞) with mean measure *c*_1_*u*_1_*x^−α^* where *α* = λ_0_/λ_1_. See the proof of (i) in Section 9 of [5].

The points in the Poisson process indicate the contributions of the various type one families to the limit *V*_1_, so if we let *x*_1_ > *x*_2_ > *x*_3_… be the points, then the jth largest family makes up a fraction *X_j_*/*V*_1_ of the population. There are three classes of mutations in the two-phase model

- type 0: Neutral mutations that occur to type 0 individuals.
- type 1A: Advantageous mutations that turn type 0 individuals into type 1.
- type 1: Neutral mutations that occur to type 1 individuals.

The type 0 mutations will have site frequency *C*_0_/*f* by the argument in Section 2.

### Theorem 4.

*The number of type 1A mutations with frequency ≥ f will be asymptotically Cf^−α^ where α* = λ_0_/λ_1_.

*Proof*. The arguments above show that the number of contributions to *V*_1_ that are larger than *x* is asymptotically *Cx^−α^*.

### Theorem 5.

*The number of type 1 mutations with frequency ≥ f will be asymptotically ν*/(λ_1_*f*).

*Proof*. We follow the derivation of Theorem 1. If we let *N*(*s*) = *Z*_1_(*s*), then the number of type 1 mutations satisfies

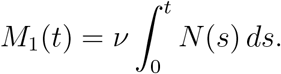

There is no λ since we assume mutations occur continuously during the life of an individual.

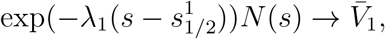

so we have 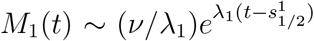. If we set 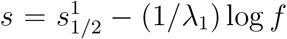, we have the desired result.

Another less intuitive approach for finding the site frequency spectrum uses the fact that the proportion of 1*A* families follows the Poisson-Dirichlet distribution PD(*α*, 0), as noted by Jason Schweinsberg. See the remark after Theorem 5 in [4]. As before, *α* = λ_0_/λ_1_. We can then apply Theorem 2 on each 1*A* family to find the site frequency spectrum for type 1 mutations. From this, we find

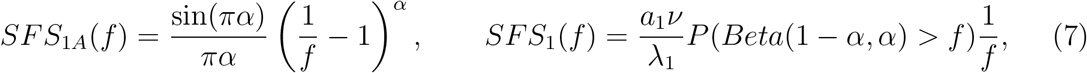

where *SFS*_1*A*_(*f*), *SFS*_1_(*f*) are the number of type 1*A* and type 1 mutations with frequency greater than *f*, respectively, and Beta(1 − *α, α*) is the Beta distribution with pdf Γ(1 − *α*)Γ(*α*)*x*^−*α*^ (1 − *x*)^*α*−1^.

To illustrate the results in Theorems 4 and 5, we turn to simulations

**Figure 1:**
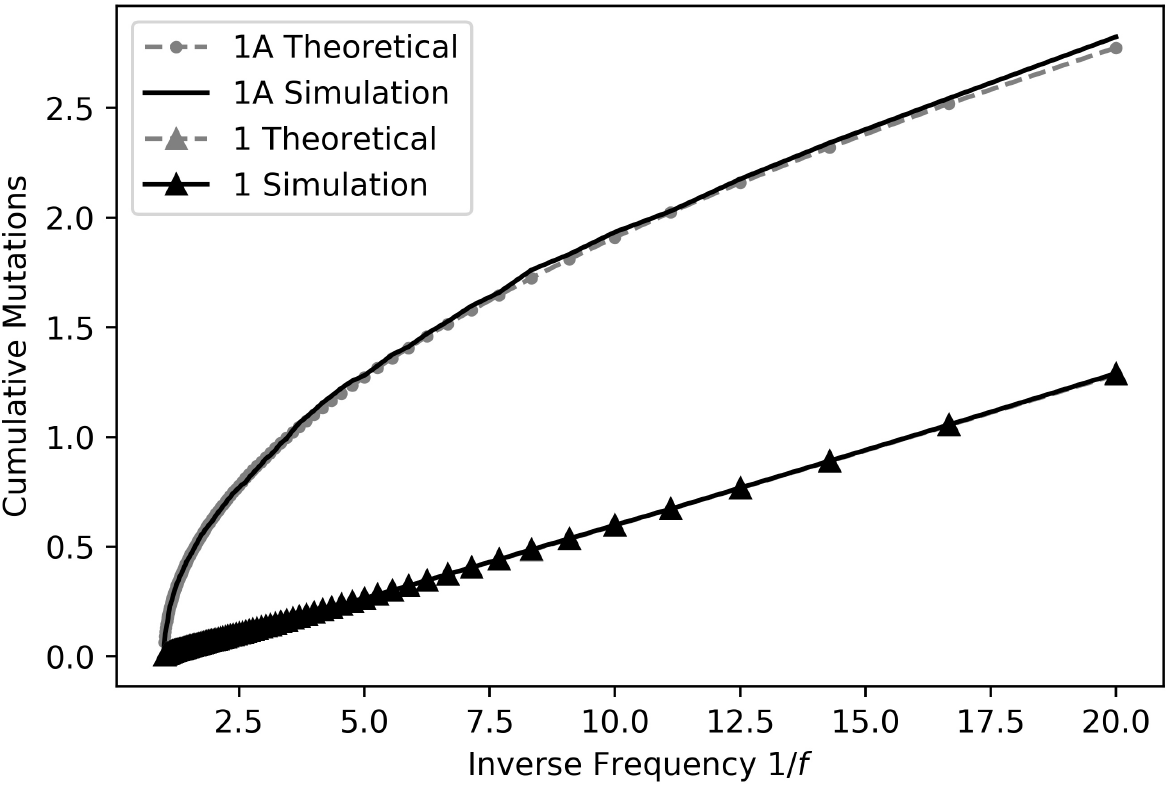
The figure shows the contribution of 1*A* and 1 families to the site frequency spectrum. The simulation was performed with parameters *ν* = 3 × 10^−3^, *u*_1_ = 3 × 10^−5^, λ_0_ = 0.02, λ_1_ = 0.04 and *a*_0_ = *a*_1_ = 1 and is the average site frequency spectrum of 1000 runs. We simulated the 1*A* families and obtained type 1 mutations for each 1*A* family by applying Theorem 2. The theoretical points are drawn from equations (7). As suggested from Theorem 5, the type 1 site frequency spectrum is linear when plotted against 1/*f*. The 1*A* lines match well and look similar to a power law, as suggested by Theorem 4.

## 4 Random fitness increases

MacDonald, Chakrabarti, and Michor [8] considered the case in which type 1 individuals have growth rates that are normal with mean μ and standard deviation *σ*. Early work on models with random fitness increases in the two-type model led to very unusual behavior in the limit t → ∞, see [6]. Results in that paper show

- If the fitness distribution was bounded then, as t → ∞, individuals with fitnesses that were close to the upper limit dominated the population.
- If the distribution was unbounded, then the population could grow faster than exponential.

In this section, we will modify our example from Section 3 so that type 1 individuals have growth rates drawn from the normal distribution with mean *μ* = 0. 04 and standard deviation *σ* = 0.005. We will see that in contrast to the limiting results just mentioned, random fitnesses do not substantially change the behavior.

The total number of type 0 individuals that have been born before time *t* is

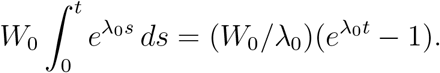

Dropping the −1, we see that when we set λ_0_ = 0.02, *t* = 1000 and take into account the mean of (*W*_0_|*W*_0_ > 0) is 50, the expected number of successful type 1 mutations is

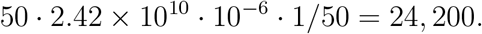

The probability that a normal is more than 4 standard deviations above its mean is 1/31,600, so the number of mutants that are 4 standard deviations above the mean is Poisson(0.766). If a few did occur, they would most likely occur close to time 1000 and hence have small family sizes.

To find the distribution of the growth rates of the mutations with the largest family sizes, we note that a mutant that occurs at time *s_i_* and has growth rate λ_1,*i*_ will grow to size *W*_1_ exp(λ_1,*i*_(1000 − *s_i_*)) at time 1000. The number of *i* that are successful and have λ_1,*i*_(1000 − *s_i_*) > *x* is Poisson with mean given by the following integral

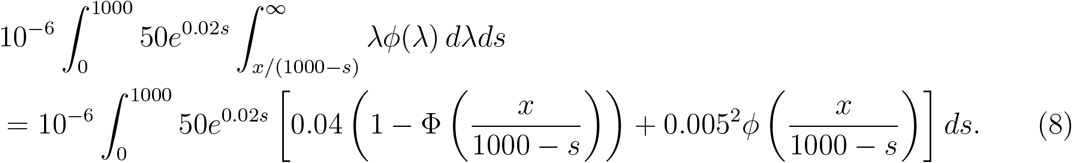

where *ϕ* and Φ are the density function and distribution function, of a normal distribution with mean *μ* = 0.04 and *σ* = 0.005. The equality follows from substituting *u* = (λ − 0.04)^2^ for the inner integral. Figure 2 graphs (8).

**Figure 2:**
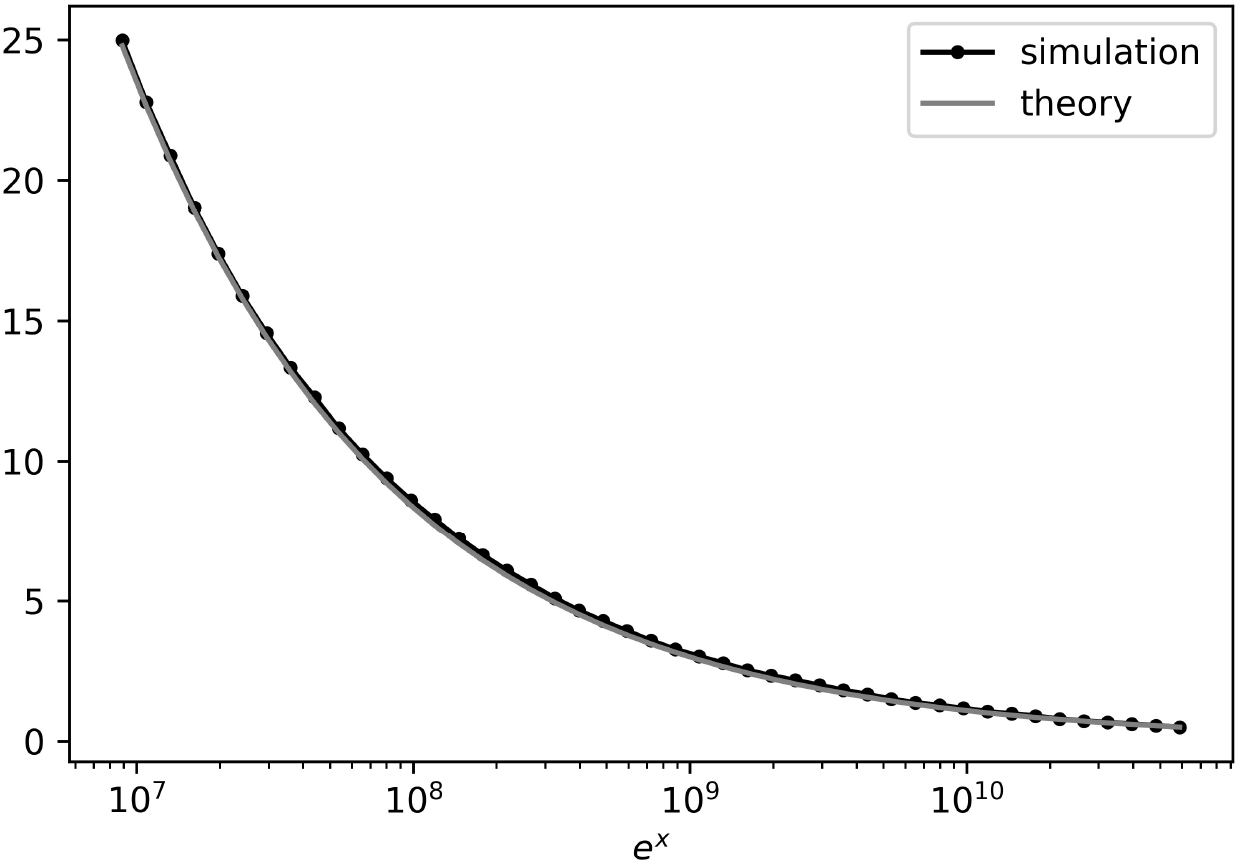
The graph indicates the expected number of 1*A* families with λ_1,*i*_(1000 − *s_i_*) > *x*. The parameters are almost the same as in (2); rather than a single λ_1_ for all type 1 families, we have a different λ_1,*i*_ for each type 1A family. Each λ_1>*i*_ is normally distributed with mean 0.04 and standard deviation 0.005. 500 runs were done up until time t = 1000. The graph shows that on average there is one family with *e^x^* > 10^10^. If the λ_1,*i*_ of the largest family is within 2 standard deviations, then multiplying *e^x^* by 1/λ_1,*i*_ implies a family of magnitude around 2 × 10^11^ or greater.

The random fitnesses cause the relative sizes of the contributions of mutations to the final population to change, but as Figure 3 shows, the site frequency still has the form *C*/*f^β^*, where *β* ≤ *α* and achieves equality in the case of non-random changes, ie *σ* = 0. The authors of [8] claim that the site frequency spectrum in the two-type model is 1/*f*. However, their simulation methods take the very crude approach of considering the binary split process until 1,000 or 1,000,000 cells are produced. This corresponds to 10 and 20 generations respectively. To make it possible for something to happen in this short amount of time the mutation rate for advantageous mutations is set to be 0.1 in the 1000 cell scenario, and to 0.03 when there are 1,000,000 cells. At birth, each cell acquires a Poisson mean 100 number of mutations. In contrast our simulations run for approximately 1000 generations, leading to populations of order 10^9^ cells, and neutral mutations occur slowly, leading to genealogical relationships that are more like those found in growing cancer tumors.

**Figure 3:**
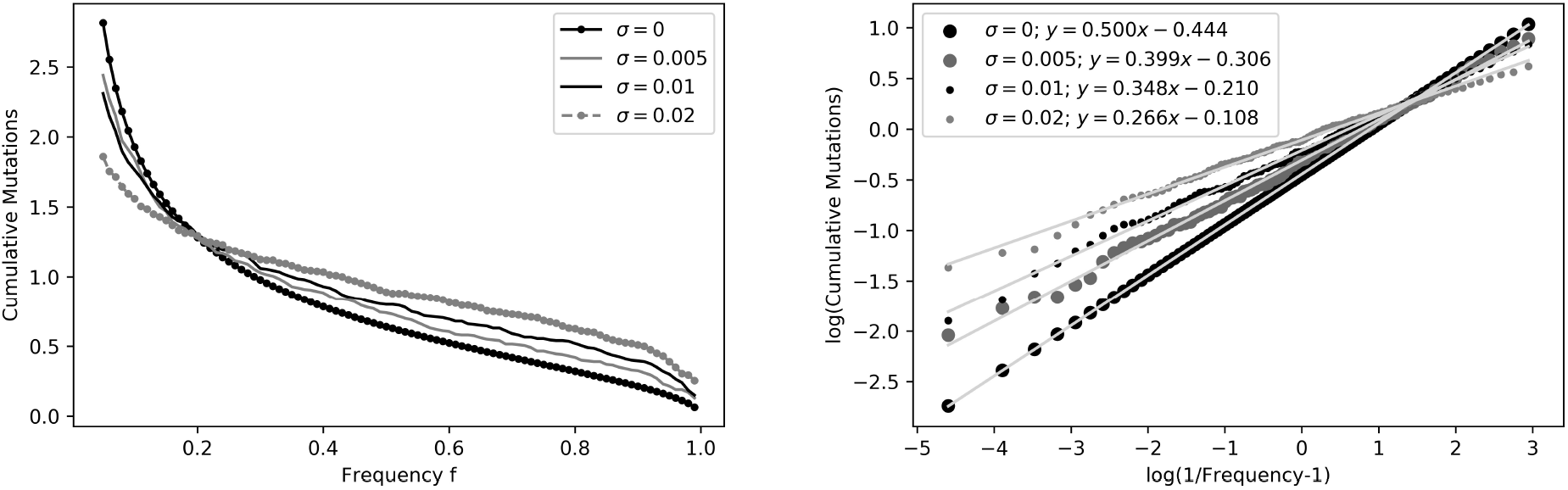
The graph on the left shows the site frequency spectrum for multiple values of *σ*. The other parameters are the same as in Figure 2. As the contribution from neutral mutations is negligible, we will only show the contribution from 1*A* families. The line for constant, ie *σ* = 0, is plotted from theory; the others are plotted from simulations with 200 runs. As *σ* increases, the expected size of the frequency of the largest mutation increases. Also, fewer mutations reach above the 0.05 frequency threshold. The graph on the right displays the same data with a log-log plot. The slopes β of the linear fits indicate that the site frequency spectrum takes the form *C/f^β^*, with β decreasing as *σ* increases.

## 5 Subclonal mutation frequencies

Bozic, Paterson, and Waclaw [3] argue that “the fact that no subclonal driver is present at intermediate frequencies cannot be taken as proof of neutral or *effectively neutral* evolution. It can be a consequence of population dynamics which create only a short window during which the driver mutation can be detected but not fixed in the population.”

To argue for this viewpoint, they use the two-phase model introduced in the Section 3 but with different notation

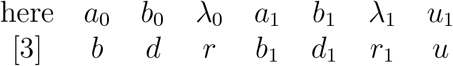

In addition they define *c* = *r*_1_ /*r* > 1, and *g* = *c* − 1. They assume that the mutation to type 1 occurs at time 0 and run the process until the time *t* at which the total population size is *M*. Let *X*_0_ be the population of type 0’s when the mutation occurs. Since *X*_0_ is large, *X_t_* ≈ *X*_0_*e^rt^*. The type 1 population at time *t* is *Y_t_* ≈ *W*_1_ *e^rct^*, where *W*_1_ is an exponentially distributed random variable with rate *cr*/*b*_1_. Note that as in Bozic et al [2] the possibility of subsequent driver mutations is ignored. As Figure 4 shows, that change is does not lead to a substantial error.

**Figure 4:**
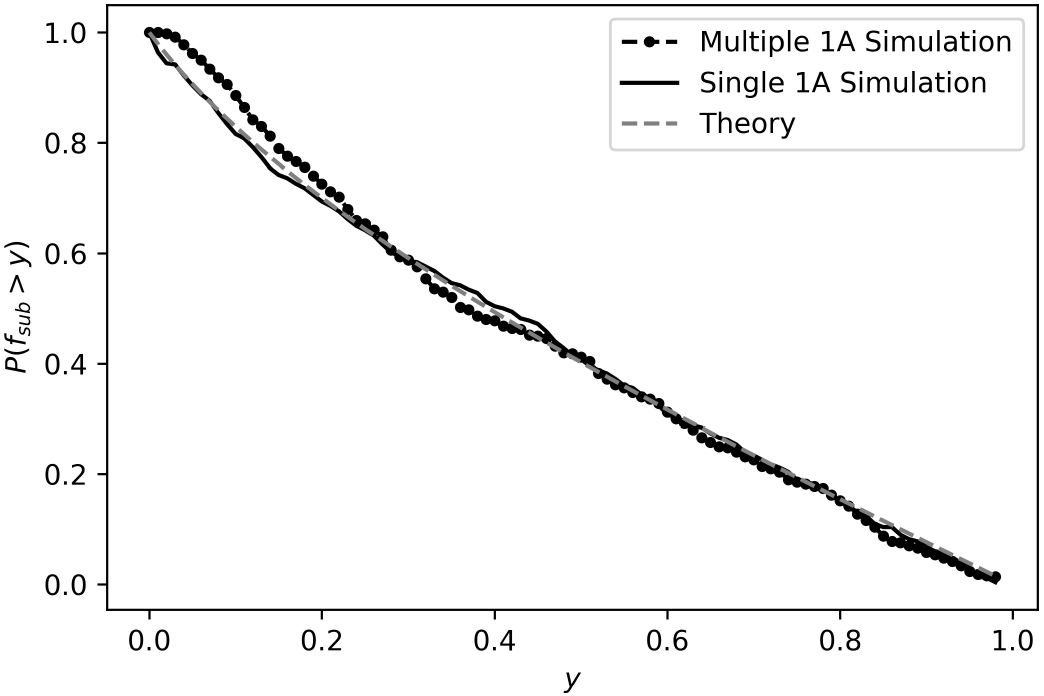
This graph gives the probability of having a driver with frequency greater than y once the tumor reaches size 10^9^. The parameters used are *a*_0_ = *a*_1_ = 1, λ_0_ = 0.02, λ_1_ = .035 and *u*_1_ = 10^−5^ and the data was generated from 500 runs. Single 1A refers to approach taken by Bozic et al. where there is only 1 selective mutation. Multiple 1A is our approach. The theory curve comes using a Riemann sum with interval size 500 to evaluate the integral in equation (9).

Writing *f_sub_* = *Y_t_*/(*X_t_* + *Y_t_*) they prove that when the total tumor size is *M* = *X_t_* + *Y_t_* the subclonal mutation frequency has

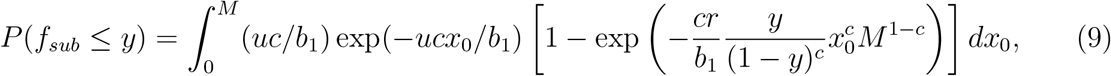

which is (1) in [3]. From this they can compute the probability of a subclonal driver being detectable, that is, *P*(0.2 ≤ *f_sub_* ≤ 0.8)

To see what this complicated formula implies, the authors turn to simulation. The mutation rate to produce an additional driver is *u* = 10^−5^. Panel a of Figure 2 shows a moderately growing tumor *b* = 0.14, *r* = 0.01, panel *b* a fast growing tumor *b* = 0.25, *r* = 0.07, and panel *c* a slowly growing tumor *b* = 0.33, *r* = 0.0013. For moderate values of selection, e.g. *g* = 30%, the probability that a driver mutation is in the detectable range [0.2,0.8] is < 15% for population sizes up to *M* = 10^9^ cells and remain below 1/3 for *M* ≤ 10^11^. For other cases considered there (*g* = 70% and 100%) the chance of detecting the subclonal driver is always < 60% and for a broad range of sizes is less than 30%.

Panels d,e,f in their Figure 2 show the frequency of a subclonal driver in the case of moderate growth when the size *M_d_* = 10^7^, *M_e_* = 5 · 10^10^ and *M_f_* = 2 · 10^8^. In the three cases the frequency is near 0, near 1, and almost uniformly distributed on [0,1]. To compare with [3] we take

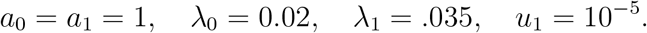

In this case, it follows from (3) that

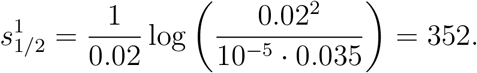

Rather than study the tumor when it reaches a fixed size, we will derive results at a fixed time by using Theorem 3. Recall that we have set 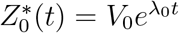 and have shown

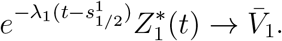

From this we see that

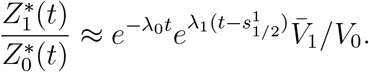

Ignoring 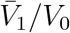 we see that

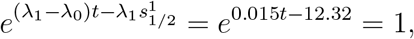

when *t* = 12.32/0.015 = 821.

It is not easy to do explicit calculations with the distribution of 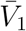 but it is easy to see that 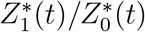 goes from 0.2/0.8 = 1/4 to 0.8/0.2 = 4 in time ln(16)/0.015 = 184, confirming that the window in which competing subclones coexist is short. It is interesting to note that we have shown that the mutations within the growing type 0 and type 1 families follow the 1/*f* law. Thus, in the single 1A approach the mere coexistence of two subclones does not ruin the 1/*f* site frequency spectrum, since there is just one 1A mutation.

## 6 Conclusions

Work of Sottoriva and Graham [11] and their co-authors [14] has shown that an exponentially growing tumor has a 1/*f* site frequency spectrum. This result has a simple derivation but the claim has drawn a large amount of criticism. Many of these concern the quality of the data used. Here, we have performed a mathematical analysis to show that the site frequency spectrum can be used to distinguish neutral evolution from one specific type of selection.

Here we have studied the two-type model of cancer evolution in which the exponentially growing population of type 0 cells can mutate to a fitter type 1, and all cells can experience neutral mutations. In this model there are three types of mutations that we call 0, 1*A*, and 1. Type 0 mutations are neutral, occur to type 0 individuals, and have a 1/*f* site frequency spectrum. Type 1 mutations are neutral, occur to type 1 individuals, and again have a 1/*f* site frequency spectrum. Type 1A mutations are selective, occur to type 0 individuals, and result in type 1 individuals. When the two types have growth rates λ_0_ < λ_1_, where *α* = λ_0_/λ_1_, then the site frequency spectrum has the shape 1/*f*^*α*^ due to 1A mutations which are more numerous than the others.

McDonald, Chakrabarti, and Michor [8] have used the two-type model to suggest that models with selection can have a 1/*f* site frequency spectrum. Our results in Section 3 show this is not true when type 1 mutations all have the same fitness increase. Their model has random increases in fitness, but in Section 4 we show that this feature does not significantly change the qualitative features of the site frequency spectrum.

Bozic, Paterson, and Waclaw [3] study the two-type model and show that it is difficult to capture a subclonal driver mutation at intermediate frequency. Their model allows only one type 1A mutation. Using our simple analytical results and computer simulations, we confirm that this prediction holds in the two type model without that restriction. However, we also show that the presence of the two subclones of comparable size in the single 1A approach does not ruin the 1/*f* site frequency spectrum. In the full model, it is the 1A mutations that change the site frequency spectrum to 1/*f^α^*.

